# Physiotherapists perceptions on informed consent and role in the healthcare system, in Europe

**DOI:** 10.1101/612473

**Authors:** Nadinne Roman, Silviu Caloian, Angela Repanovici, Roxana Miclaus, Gabriela Sechel, Liliana Rogozea

## Abstract

**Introduction:** Physiotherapy has developed over the last century, and the physiotherapists’ professional identity is growing. The heterogeneity of physiotherapy studies in Europe, local government, and health policies have influenced the responsibilities and ethical reasoning of physiotherapists. Our study aims to explore the perceptions and differences regarding informed consent (IC) and the role of physiotherapists in healthcare in an educational, legislative, and health policy context.

**Material and Methods:** A cross-sectional survey was distributed online to physiotherapist graduates in Europe. The survey contained two open questions regarding IC and assumed role in healthcare. The data was operated to and analyzed by using a theory-based approach (open and axial coding), providing a qualitative spectrum of categories for the two items linked on IC and the role in healthcare.

**Results:** Eight categories of issues related to IC and seven categories related to the role in healthcare were identified. The physiotherapist graduates from Romania, France, Belgium, Italy, and other countries from inside and outside European Union response rate was 81.85% (n = 248 from 303) for the item related to IC and 71.62% (n = 217) to the second item related to the role in healthcare. A percent of 24.20% (n=60) are still considering IC a simple patient information process, while 23.40% (58) have linked IC with ethically and legally issues, 21.67% (n=51) of physiotherapists are minimizing their role in healthcare at restoring physical independence, while 6.91% (n=27) are aware of their multidisciplinary role. The country comparison analysis revealed that physiotherapists from UK and Italy are more aware regarding IC and that physiotherapists from Belgium and France are better oriented regarding their role in healthcare.

**Conclusions:** The study shows that heterogeneity, legislation, and healthcare system differences influence physiotherapists professional development. Future research is needed to establish the reason for the reduced perception of physiotherapists regarding their role as health promoters.

## Introduction

Throughout the evolution of the physiotherapist profession, the concept of professional identity has developed, assuming the existence/presence of a set of moral norms, abilities, and skills designed to define the profession of the physiotherapist towards high standards (1)(2)(3). The occupation of the physiotherapist has developed throughout Europe during the last century as a profession assimilated to healthcare professionals (4)(5)(6). In most countries in Europe and around the world, the profession of physiotherapy has been regulated and recognized so far, leading to the occurrence of ethical norms that must be respected and compliance with high standards of practice (7)(8)(9). Romania is the only country of the European Union where the physiotherapist profession was partially regulated in 2016, and the professional organization has incomplete legislation (10).

The professional identity of those performing social work, as in the case of physiotherapists, is a continuous process that is determined by the context, factors, or situations that may occur at work. It is linked to specific roles that are attributed more from a legal point of view than relative to the complexity and potential conceptualization of each profession. Formation of the professional identity of physiotherapists takes place over time. The workplace, the staff involved in everyday activities, the moral principles and values of each, the accumulated experience, and the moral conflicts arising from the intersection of individual values and beliefs are elements which influence the accumulation of professional skills (11)(12).

The legislation, health systems, and study programs in physiotherapy vary from one country to another, although competencies are mostly similar, with few exceptions. However, the differences between countries health systems are essential, especially in the context of migratory physiotherapists, especially in European territory. According to the European database on regulated professions, from 1997 to 2013, 19 973 physiotherapists migrated to European countries. Ethical issues related to the profession of physiotherapy also arise due to different systems and health legislation in the context of migration (13) (14)(15).

In addition to the physical and functional rehabilitation skills and abilities, physiotherapists play a vital role in the prophylaxis, health, promotion of physical activity and long-term outcomes after different surgical procedures, being part of the medical profession with a significant impact on the quality of life (16, 17).

The ethical issues encountered in medical practice, legislative norms, research, and medical technology, marks this multidisciplinary profession, which influences medical practice and population. In the context of the heterogeneity of studies, legislation, health systems, and the needs of the high population, physiotherapists must provide high-quality health services and be involved in promoting health. Considering the increased number of elderly and disabled people, as well as the risk of cardiovascular disease and premature death due to physical inactivity worldwide, physiotherapists have to respect high professional standards (18)(19)(20).

So far, research shows that the professionalism and attitude of the physiotherapist have considerable repercussions on the evolution of the rehabilitation process (21)(22). As a medical act in which the physiotherapist often guides the patient in the physical therapy program, the approach of physiotherapists is often paternalist and often does not involve the patient in creating the content of the physiotherapy program, omitting cultural or social elements of patients, and often granting importance only to physical rehabilitation and less motivation through socio-cultural aspects (23)(24).

Additionally to the ethical issues encountered in medical practice, the current legislative norms, as well as the evolution of medical research and technology, make their mark on this multidisciplinary profession, resulting in multiple professional responsibilities affecting the quality of healthcare services. Considering the specificity of physiotherapy care, which involves many issues, create difficult circumstances for both patients and physiotherapists and medical staff. Physically and emotionally prolonged contact of the patient with the physiotherapist, the lack of a specific framework for obtaining IC, but also the legislative, educational and professional differences specific to each country influence the physiotherapy practice. The ethical issues encountered in the physiotherapist’s medical practice are also difficulties of practice. (25)(26)(27)(28)(29).

Regarding both patients’ and physiotherapists’ perspectives about the rehabilitation process, the most important ability appreciated by patients is communication, followed by the therapist’s focus on the patient himself and not just on the present disorder. These aspects confirm the need to develop a professional relationship between the patient and the therapist, precisely because of the primary purpose of rehabilitation, namely the patient’s autonomy. Within the relationship developed with the patient, the therapist should draw some guidelines and recognize critical ethical issues, but also extract ethical principles and connect them to particular clinical contexts (31). It is crucial for the physical therapist to assume the level of practical authority, to have the ability to connect ideas, to understand the relationships and to have the knowledge necessary to make the right decision finally.

Regarding the process of obtaining informed consent (IC), one of the particularities of applying the therapies in physiotherapy, but also from the perspective of obtaining informed consent, is related to the specificity and dynamics of the physical therapy sessions. In most cases, techniques and methods used throughout treatment change, often even during a physiotherapy session, so the physiotherapist has to make multiple decisions during a treatment session. There are no strict protocols, and such protocols cannot be applied, and if they exist, physiotherapy is a medical branch that treats the patient 90% and less the illness itself. The previous studies regarding IC show that most physiotherapists obtain patient consent at the onset of treatment, along with the patient evaluation, not repeating this process in every change of techniques or maneuvers used (32)(24)(28).

Given the many factors that may influence the physiotherapists’ professionalism, the main objective of our research was to explore physiotherapists’ perceptions of informed consent (IC) and the level of development of professional identity in the context of educational, legislative and professional differences.

## Material and methods

To investigate the perception of physiotherapists in the medical practice, we have sought to contextualize the concepts and ethical principles resulting from the literature revision. A descriptive, qualitative and comparative study was carried out through a questionnaire survey. The operationalization of the ethical elements extracted through the review of the literature was carried out to link the ethical concepts and premises of reality, to be able to objectively measure the perception of physiotherapists on ethical aspects of professional practice. Through the process of operationalization, we identified several dimensions of ethical issues in the physiotherapy care process, which we later transposed into a questionnaire addressed to physiotherapists.

We have distributed a questionnaire addressed to graduate physiotherapists among several European Union countries. The questionnaire used was qualitative research in a small percentage, with two open questions about physiotherapists’ perception of IC and their role in healthcare.

The survey was distributed online through the Survey Monkey platform. Before applying the survey, we have obtained the approval of The Ethics Commission of the Faculty of Medicine of the Transilvania University of Braşov. We have disseminated the questionnaire online, by contacting different professional physiotherapy associations within the European Union, and as well through the social media pages of professional groups. The questionnaires were distributed from November 2017 until July 2018, initially in Romanian, and after were translated into English, French, and Italian, by translators of specific languages and then distributed once again. The sampling was random.

The sample size was 303 respondents. The questionnaire was distributed to physiotherapists in an online environment using physiotherapy groups from the social networks in Romania, Belgium, France, the United Kingdom, Spain, Portugal, Switzerland, Hungary, Italy and other groups without specific geographical allocations. We have contacted all physiotherapy associations in Europe, but most of the answers were negative or the response was lacking. The informed consent was obtained implicitly by filling in the questionnaire. No personal data has been collected under European law. Initially, 331 questionnaires were collected, of which 28 were removed due to missing or incomplete data. In the case of the two open-ended questions, the answer rate was 81.85% (n = 248) to the first open question related to CI and 71.62% (n = 217) to the second item related to the role in healthcare.

## Data operationalization

### Informed Consent

The responses gathered from the two open items varied and it was necessary to distinguish between the types of response. Depending on the description, several categories have been differentiated by operating and encoding data. Inductive coding was applied separately by two of the team members, following a few steps: 1) Reading raw data and creating codes to cover the sample based on responses; 2) re-evaluating the sample and applying the codes for each respondent and creating another code for irrelevant answers that did not correspond to the question. We have added a code for respondents which claimed the lack of IC at the onset of physiotherapy. A third member of the team has resolved misunderstandings.

After the coding, the responses were separated into seven categories, making an ordinal grading, accounting as maximum complexity of responses related to patient rights and legal considerations, and at the last level classifying the respondents who stated that they do not demand IC at the onset of treatment.

The coding of responses was done both to obtain data that can be measured and analyzed and to be ranked according to the efficiency or the correctness of the response, allowing us to order them on an ordinary scale:

0 = assigned to respondents who do not request IC,

1 = assigned to irrelevant answers,

2 = assigned to respondents who believe that obtaining IC is done to protect themselves against malpractice

3 = assigned to respondents who declared it to be an information process about the therapeutic program

4 = assigned to respondents who thought IC is necessary to increase patient confidence

5 = assigned to respondents who sought IC as a tool to determine good collaboration with the patient and to integrate it into the physical therapy process

6 = assigned to respondents who have declared patient consent or disagreement with the treatment

7 = the most comprehensive and correct answers from legal, ethical, and professional perspectives.

### Assumed role in healthcare

The second open question item of the questionnaire aimed to investigate the perception of physiotherapists role in healthcare, as medical care providers. Data interpretation included a pre-review of all responses and their categorization. Inductive coding was applied separately by two of the team members, following a few steps: 1) Reading raw data and creating codes to cover the sample based on responses; 2) re-evaluating the sample and applying the codes for each respondent and creating another code for irrelevant answers that do not correspond to the question. A code added for those who said they did not ask for IC at the onset of physiotherapy. A third member of the team has resolved misunderstandings. This item of the questionnaire had a response rate of 71.62% (217 out of 303).

Response categories were differentiated into eight sections. The most complex and appropriate category was considered the last one. The responses related to the prophylactic, therapeutic, and rehabilitation role, which is, in fact, the definition of physiotherapy (16) and embraces the complex characteristics of the physiotherapist profession.

0-Irrelevant answer

1-Role in rehabilitation

2-Restore physical / independent condition

3-Planning, evaluation, and application of physiotherapy

4-Improving the quality of life/condition

5-Interdisciplinary / binder in the medical team

6 Essential / Complex

7-Prophylactic, curative and therapeutic

## Results

### Demographic Analysis

Statistical analysis was performed using SPSS 20. Descriptive analysis of the two items was accomplished, and for comparative analysis, the non-parametric Kruskal-Wallis test was used to determine the differences perceived related to IC and role in healthcare according to age, experience, type of practice and countries of origin. Subsequently, pairwise comparisons were performed using Dunn’s (1964) procedure with a Bonferroni correction for multiple comparisons. Adjusted p-values are presented.

### Distribution by type of practice

Within the first item, out of 248 respondents, 145 (58.47%) respondents came from the private sector, 94 (37.91%) from the public sector and 9 (3.62%) from other mixed sectors. In the second item, out of 217 respondents, 125 (57.61%) came from the private sector, 85 (39.18%) and 7 (3.21%) in the mixed sectors.

### Descriptive analysis

## Discussions

### Informed Consent

The results of the statistical analysis revealed that the physiotherapists from Italy and UK had been mostly focused on the complexity of the IC process so that 19 of 36 and 13 of 21 (Table 2) respectively provided adequate responses to the patient’s consent with the treatment plan and the ethical aspects and legal IC. The results are surprising given the differences and similarities between the analyzed groups. Thus, as in Belgium and UK, the physiotherapists have professional independence, with diagnostic attributions as autonomous practitioners, whereas in France, Italy, and Romania, physiotherapists have a secondary contact with a patient at the referral of a specialist physician and have no competence of diagnostic (33). Previous literature often advocates the need for a concise framework for IC in physiotherapy, given the specificity and dynamics of treatment in this medical division (34)(35).

**Table 1:** Age and experience distribution

| Age | Item 1<br>number/percent | Item 2<br>number/percent | Experience | Item 1<br>number/percent | Item 2<br>number/percent |
| --- | --- | --- | --- | --- | --- |
| 21-30 Years | 105/42.33 % | 94/43.31% | 1-3 Years | 77/31.05% | 65/29.96 |
| 31-40 Years | 100/40.32 % | 85/39.17% | 3-5 Years | 29/11.69% | 31/14.29 |
| 41-50 Years | 33/ 13.30 % | 27/12.44% | 5-10 Years | 60/24.20% | 51/23.50 |
| 51-60 Years | 7/ 2.82 % | 8/3.68% | 10-20 Years | 53/21.37% | 46/21.19 |
| > 60 Years | 3/1.20 % | 3/1.38% | >20 Years | 29/11.69% | 24/11.06 |
| Total | 248/100.00% | 217/ | Total | 248/100% | 217/100.00% |

**Table 2.** Crosstabulation of reason for IC demanding at physiotherapy onset by countries respondents

| Count | Reason for IC demanding * Countries Crosstabulation |  |  |  |  |  |  | Total |  |
| --- | --- | --- | --- | --- | --- | --- | --- | --- | --- |
|  | Country distribution |  |  |  |  |  |  | N | Percent |
|  | Romania | Italy | France | Belgium | UK | UE | Non-UE |  |  |
| Do not demand | 5 | 1 | 0 | 0 | 0 | 0 | 0 | 6 | 2.41% |
| Irrelevant answers | 10 | 1 | 0 | 2 | 0 | 0 | 2 | 15 | 6.04% |
| Physiotherapist protection/<br>malpractice | 6 | 3 | 6 | 2 | 0 | 0 | 6 | 23 | 9.27% |
| Patient information | 40 | 3 | 5 | 4 | 3 | 2 | 3 | 60 | 24.20% |
| To gain patient's trust | 7 | 0 | 0 | 0 | 0 | 0 | 2 | 9 | 3.63% |
| Collaboration with patient | 14 | 3 | 6 | 5 | 0 | 0 | 1 | 29 | 11.70% |
| For patient Consent | 18 | 6 | 4 | 3 | 5 | 3 | 9 | 48 | 19.35% |
| Ethical and legal issues | 11 | 19 | 1 | 4 | 13 | 2 | 8 | 58 | 23.40% |
| Total | 111 | 36 | 22 | 20 | 21 | 7 | 31 | 248 | 100.00% |
| Percent | 44.75% | 14.52% | 8.89% | 8.07% | 8.46% | 2.82% | 12.50% | 248 | 100.00% |
| Mean | 4.01 | 6.04 | 4.11 | 4.24 | 6.15 | 5.17 | 4.75 |  |  |
| Sd | 1.942 | 1.870 | 1.711 | 1.895 | 1.424 | 1.722 | 2.111 |  |  |
| 95% CI Lower Bound | 3.61 | 5.23 | 3.26 | 3.21 | 5.48 | 3.36 | 3.86 |  |  |
| 95% CI Upper Bound | 4.41 | 6.85 | 4.96 | 5.26 | 6.82 | 6.97 | 5.64 |  |  |
| Minimum | 0 | 0 | 2 | 1 | 3 | 3 | 1 |  |  |
| Maximum | 7 | 7 | 7 | 7 | 7 | 7 | 7 |  |  |

Although the need to obtain IC from the patient is found in national legislation, both in Romania and in other countries, there are probably several factors influencing the process of requesting and obtaining IC from patients. Five Romanian physiotherapists claimed the lack of IC and 36.03% associated this process by providing treatment information. In Romania, the heterogeneity of university studies, the lack of a professional association and a code of conduct for physiotherapists are elements that might influence this process. Also, in France, it is likely that the influence of this legal and ethical aspect is related to the type of university studies provided by the French higher education system. Starting in 2015, the reform of masso-kinesitherapy studies is taking place, extending for another two years, and reaching five years of university studies (36). At the European level, are voices that challenge the skills and abilities of physiotherapists in specific countries across the EU that say there is no uniformity in physiotherapy studies across the European Union. The Secretary-General of the Maso Therapy Order in France states that Belgium and France are the most demanding in terms of studies, and the weakest and most critical educational systems reported in the physiotherapist profession are those in Romania and Spain. (37). Our results show that French and Belgian physiotherapists do not have a complex perception regarding IC and ethical aspects of this process are minimized.

The results obtained reveal that of the 248 respondents, 24.20% (Table 2) linked the IC process to simple patient information, s omitting the ethical and legal issues that were created and implemented as practice standards, continuing to use the IC process as a tool for detailing the treatment objectives and the techniques used (38)(24).

An important aspect to consider is professional autonomy and initial contact with the patient. Considering the legislative aspect, specific to each country, the physiotherapists performing the medical activity, only at the physician’s indication, are having secondary contact with the patient, so the ethical and professional responsibilities lie primarily with the physician and less with the physiotherapist. We cannot extrapolate on this, because in Italy for example, where the situation is similar to Romania and physiotherapists work under the guidance of a physician, 52.80% (19) of the respondents responded appropriately to this item. Italian physiotherapists considering that the reason for requesting and obtaining IC is represented by the ethical, deontological, and legal nature of the profession and the process itself.

Differentiating response categories by the reason physiotherapists believe that IC is obtained has allowed us to point out important aspects of physiotherapists’ perception of IC. So far, this aspect has not been studied from this perspective, and the responses suggest that the IC process is elaborate. In addition to legal and deontological considerations, obtaining IC is a useful tool that allows physiotherapists to establish a connection and a relationship with the patient. (32) (39)

It seems that the lack of a concrete framework for IC obtaining in physiotherapy, due to treatment specific and dynamics are negatively influencing the perceptions of physiotherapists regarding IC even after a century of professional development (6)(38).

### Assumed role in healthcare

From the summary analysis of the results, it is observed that the highest percentage was attributed to the role in the physical rehabilitation process (21.67%) (Table 3), omitting the other professional competences. It should be mentioned that the “Interdisciplinary” role was representative of the Romanian participants - 7 out of 10 responses. We want to emphasize this aspect because of the interdisciplinary relationship between Romanian physiotherapists and specialists, which presupposes a rather collaborative relationship between the two and not a hierarchy, although as discussed previously, the physiotherapist cannot apply physiotherapy techniques and methods without the referral of the physician. (40)

**Table 3.** Cross-tabulation of physiotherapists role in healthcare by countries respondents

| **Assumed role in healthcare \* Countries Crosstabulation** |  |  |  |  |  |  |  |  |  |
| --- | --- | --- | --- | --- | --- | --- | --- | --- | --- |
| Count | Country distribution |  |  |  |  |  |  | Total | Percent |
|  | Romania | Italia | France | Belgium | UK | UE | Non-UE |  |  |
| Other | 12 | 1 | 0 | 0 | 0 | 1 | 1 | 15 | 23.50% |
| Role in rehabilitation | 24 | 8 | 4 | 3 | 7 | 2 | 3 | 51 | 21.67% |
| Restore Physical Function/<br>Independence | 21 | 6 | 2 | 5 | 6 | 1 | 6 | 47 | 11.06% |
| Planning, assessment and<br>application of physiotherapy | 13 | 0 | 2 | 0 | 3 | 0 | 6 | 24 | 10.60% |
| Improving the quality of life /<br>condition | 11 | 0 | 4 | 1 | 2 | 2 | 3 | 23 | 4.61% |
| Interdisciplinary | 7 | 1 | 1 | 0 | 0 | 0 | 1 | 10 | 9.21% |
| Essential / Complex | 9 | 4 | 3 | 2 | 1 | 0 | 1 | 20 | 12.44% |
| Promotion, prevention,<br>treatment/intervention,<br>habilitation and rehabilitation. | 5 | 7 | 3 | 8 | 1 | 0 | 3 | 27 | 6.91% |
| Total | 102 | 27 | 19 | 19 | 20 | 6 | 24 | 217 | 100.0 % |
| Percent | 47.00% | 12.44% | 8.75% | 8.75% | 9.21% | 2.75% | 11.05% | 217 | 100% |
| Mean | 2.60 | 3.74 | 3.89 | 4.18 | 2.45 | 2.00 | 3.21 |  |  |
| Sd | 1.951 | 2.734 | 2.246 | 2.604 | 1.701 | 1.673 | 1.978 |  |  |
| 95% CI Lower Bound | 2.20 | 2.56 | 2.77 | 2.84 | 1.65 | 0.24 | 2.37 |  |  |
| 95% CI Upper Bound | 3.00 | 4.92 | 5.01 | 5.52 | 3.25 | 3.76 | 4.04 |  |  |
| Minimum | 0 | 0 | 1 | 1 | 1 | 0 | 0 |  |  |
| Maximum | 7 | 7 | 7 | 7 | 7 | 4 | 7 |  |  |

Most of the physiotherapists summarize their role in the healthcare process, at the level of restoring the patient’s functional capacity and the regaining of physical autonomy, although the role of physiotherapists in healthcare is complex. The World Confederation of Physiotherapists (WCPT) states: “Physiotherapists offer services that develop, maintain and restore the maximum ability for movement and functional capacity of people. They can help people at any stage of life when movement and functionality are threatened by aging, injury, illness, disorder, condition, or environmental factors. Physiotherapists help people maximize their quality of life, with regard to physical, psychological, emotional and social well-being. They work in the health spheres of promotion, prevention, treatment/intervention, rehabilitation and provision of services that ensure maximum independence in everyday life activities through education, education and/or treatment (16). Physiotherapists from Italy and Belgium seemed to orient themselves in a higher percentage with the complexity of the profession.

Most of the WCPT physiotherapists attributions are found in the categories of responses differentiated by the present study (Table 3), but the proportion of physiotherapists that perceive their complex role in healthcare is deficient. Although it is a profession that has been developing for more than 100 years in Europe, the professional identity of physiotherapists is still in a complicated process, dynamic and with various influences (1)(3)(2).

The transition from student to practitioner, as well as the many elements that can influence the professional development of physiotherapists profession. Graduation country, workplace, experience, continuous professional training, ethical and legislative education, physiotherapist integration into the medical team, lack of autonomy and the promotion of physiotherapy as an essential medical division in patients ‘health are elements that can influence physiotherapists’ perception of their role in the healthcare system (41) (42)

### Comparative analysis

One of the issues investigated was the influence of factors such as age, work experience and country of origin in the formation of professional identity and IC perceptions.

As far as the professional identity of physiotherapists is concerned, they are still in development and they seem to have difficulties in terms of professional strength and integration into the medical team (43)(44).

The results of our research (Table 4) show that there are no differences in the perception of the role in the health system or on the ethical aspects of IC based on age or work experience. In contrast, there was a difference in terms of IC issues and the public or private sector of activity. Physiotherapists performing public health work are thus more aware of the complexity of the IC process (Table 4).

**Table 4.**
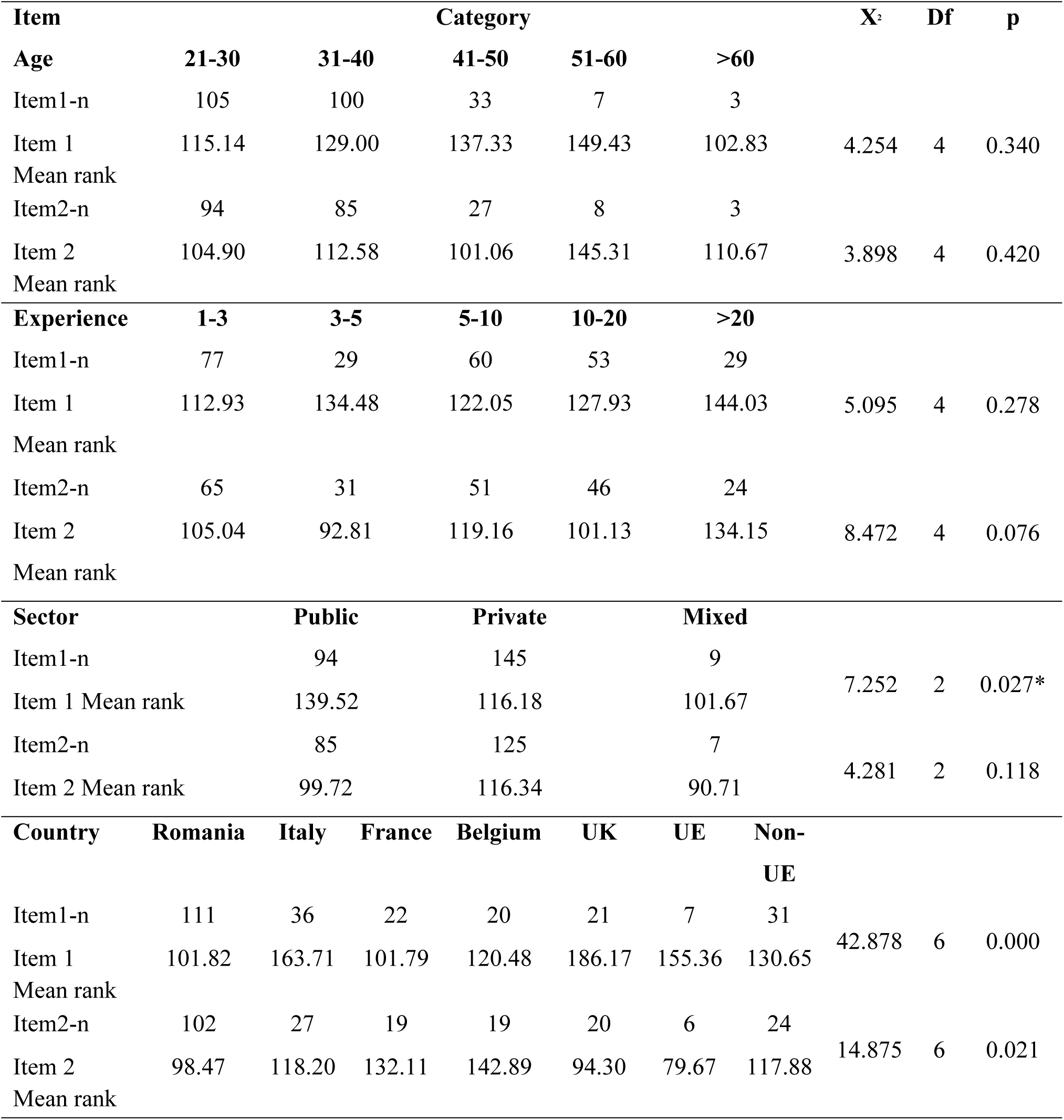
Kruskal-Wallis Independent sample test comparison by age, experience, working sector and countries

In the comparative analysis on countries (Table 4), there is a statistically significant difference in the perception of IC and professional identity, to the disadvantage of physiotherapists in Romania, France, and Belgium, versus physiotherapists in the UK and Italy. The differences of physiotherapy programs available in many countries, mainly linked to the profile of the faculties: physical education or healthcare are elements that influence the formation of physiotherapists professional identity and professional ethics, including in the UK. (45).

Also, the importance of continuing professional education has proven to be an essential factor in the development of professional identity. Moreover, we want to emphasize that physiotherapists in Romania do not currently have an obligation to attend continuous professional training (46) (47).

Pair comparison with Bonferroni correction of the Kruskal Wallis test (Table 5) denotes differences of perception both on IC aspects, but also on the role in the healthcare system and the formation of professional identity, to the benefit of physiotherapists in the UK and Italy, to the detriment of physiotherapists in Romania, France, and Belgium. In Romania, physiotherapy program studies can be either in a university with a medical profile or in a sports university. Also, in France and Belgium, this specialization is also earned from a university study program with a physical education background. Contact with the clinical component of the physiotherapist students is a factor affecting the professional training process, which takes shape during the physiotherapy studies (48)(49)(50)(51).

**Table 5:** Bonferroni correction for multiple comparisons of Kruskal Wallis test.

| Group Versus (VS) |  | Mean Ranks | Test | Std. | Std. test | p |
| --- | --- | --- | --- | --- | --- | --- |
|  |  | VS | statistic | error | statistic |  |
| Item 1 | France-Italy | 101.09/163.71 | 62.617 | 19.052 | 3.287 | 0.001 |
|  | France UK | 101.09/186.17 | -85.076 | 21.478 | -3.961 | 0.000 |
|  | Romania-Italy | 101.82/163.71 | -61.889 | 13.503 | -4.583 | 0.000 |
|  | Romania-UK | 101.82/186.17 | -84.347 | 16.753 | -5.035 | 0.000 |
|  | Romania-Non-UE | 101.82/130.65 | -28.826 | 14.301 | -2.016 | 0.044 |
|  | Belgium-Italy | 120.48/163.71 | 43.233 | 19.634 | 2.202 | 0.028 |
|  | Belgium-UK | 120.48/186.18 | -65.692 | 21.996 | -2.987 | 0.003 |
|  | Non-UE-UK | 130.65/186.17 | 55.522 | 19.897 | 2.790 | 0.005 |
| Item 2 | UE-Belgium | 79.67/142.89 | 63.228 | 28.976 | 2.182 | 0.029 |
|  | UK-Belgium | 94.30/142.89 | 48.595 | 19.823 | 2.451 | 0.014 |
|  | Romania-France | 98/47/132.11 | -33.640 | 15.461 | -2.176 | 0.030 |
|  | Romania-Belgium | 98.47/142.89 | -44.429 | 15.461 | -2.874 | 0.004 |

\*Pairwise comparison results regarding Public versus Private Sector on Item1: Test statistic: 23.342, std error: 9.322, Std. Test statistic: 2.504 and p=0.037

## Conclusions

The development of the professional identity of physiotherapists in Europe differs according to the educational system and the present legislation. In countries where physiotherapy programs are belonging to the medical sphere, physiotherapists have a greater sense of their professional role in healthcare, as medical services providers. The heterogeneity of university studies, their duration, the presence of professional organizations, and legislation are factors that influence the perception and attitude of physiotherapists on the IC process, both from a medical, ethical, and legal point of view.

The lack of a concrete framework for obtaining IC in physiotherapy is still felt in medical practice, and the lack of professional activity of physiotherapists focused on prevention and prophylaxis is poorly identified by them.

We recognize the limits of research through the small sample size and the inequality of the analyzed groups, and we suggest continuing the research from this perspective to determine real factors which require changes for the professional practice of physiotherapists at the European standards in all EU countries.

